# Resolving the pharmacological redox-sensitivity of SARS-CoV-2 PLpro in drug repurposing screening enabled identification of the competitive GRL-0617 binding site inhibitor CPI-169

**DOI:** 10.1101/2023.10.11.561987

**Authors:** Maria Kuzikov, Stefano Morasso, Jeanette Reinshagen, Markus Wolf, Vittoria Monaco, Flora Cozzolino, Simona Golič Grdadolnik, Primož Šket, Janez Plavec, Daniela Iaconis, Vincenzo Summa, Francesca Esposito, Enzo Tramontano, Maria Monti, Andrea R. Beccari, Björn Windshügel, Philip Gribbon, Paola Storici, Andrea Zaliani

## Abstract

The SARS CoV-2 Papain-Like protease has multiple roles in the viral replication cycle, related to both its polypeptide cleavage function and its capacity to antagonize host immune response. Targeting PLpro function is recognized as a promising mechanism to modulate viral replication whilst supporting host immune responses. However, development of PLpro specific inhibitors remains challenging. Upcoming studies revealed the limitation of reported inhibitors by profiling them through a pipeline of enzymatic, binding and cellular activity assays showing unspecific activity. GRL-0617 remained the only validated molecule with demonstrated anti-viral activity in cells. In this study we refer to the pitfalls of redox-sensitivity of PLpro. Using a screening-based approach to identify inhibitors of PLpro proteolytic activity, we made extensive efforts to validate the active compounds over a range of conditions and readouts, emphasising the need for comprehensive orthogonal data when profiling putative PLpro inhibitors. The remaining active compound CPI-169, showed to compete with GRL-0617 in NMR-based experiments, suggesting to share a similar binding mode, opening novel design opportunities for further developments as antiviral agents.

**Author summary:** The increasing knowledge about SARS-CoV-2 allowed the development of multiple strategies to contain the spread of COVID-19 infection. Nevertheless, effective antiviral pharmacological treatments are still rare and viral evolution allowed a fast adaptation and escape from available containment methods. The papain like protease (PLpro) has now become the next most promising SARS-CoV-2 therapeutic due to its multiple functions in virus replication cycle and antagonization of host immune response. However, due to inherent flexibility and sensitivity of this enzyme specific inhibitors are rare. Here we report on a screening strategy using repurposing of known drugs that takes into account PLpro characteristics to identify new inhibitors, showing the success of the approach by identifying CPI-169 that competitive targets the well described GRL-0617 inhibitor binding pocket of PLpro and helping to design further antiviral agents.

## Introduction

After over three years of pandemic SARS-CoV-2 still represents a severe threat to worldwide health and economic systems. The development of multiple vaccines against SARS-CoV-2, has improved the outcomes for infected individuals. In addition, small molecules have been brought to the clinic, including the repurposed compounds remdesivir which inhibit the RNA-dependent-RNA-polymerase (RdRp), and the main protease (Mpro) inhibitor nirmatrelvir (PF-07321332) that is approved for application in combination with ritonavir. Being key enzymes of SARS-CoV-2 replication, RdRp and Mpro were first targets for development of new specific antivirals. The papain like protease (PLpro) has now become the next most promising SARS-CoV-2 therapeutic due to its multiple functions in maturation of viral polyproteins and antagonization of host immune response. PLpro is responsible for release of nsp1-4 from the viral polyprotein and, in addition, it antagonizes host cellular ubiquitination and ISGylation processes, both playing important roles in the regulation of innate immune responses to viral infections [1–3].

PLpro is part of nsp3, the largest non-structural protein within the SARS-CoV-2 genome comprising multiple functions and functionally independent subunits within one protein [1]. SARS-CoV and SARS-CoV-2 PLpro show an overall 83% sequence identity, being highly conserved within the coronavirus family [PMCID: PMC7430346], and raising the possibility of development of a broad spectrum anti-coronaviridae inhibitors [4]. Nsp3 is composed of five domains, named from nsp3a to 3e, connected through different linkers and followed by two transmembrane regions (TM) and a Y-domain at the C-term [5]. The proteolytically active papain-like protease domain (PLpro) is part of the nsp3d region. PLpro is a cysteine protease with a catalytic triad formed by Cys111–His272–Asp286, which shows recognition preferences for LXGG↓XX motif (arrow indicates the cleavage site) [1,2]. S1, S2 form a narrow tunnel responsible for the recognition of two Glycines. S4 recognizes the hydrophobic side chains of Leu (or Ile). S3 lacks specific residue preference due to recognition of the peptide backbone [6]. Besides, PLpro has two ubiquitin binding sites (Ub1 and Ub2) located distal to the active site. Whilst SARS-CoV-2 PLpro prefers to bind ISG15, SARS-CoV PLpro preferentially cleaves the K48-linked di-Ub chains [7,8]. C-terminal from PLpro lays nsp3e that contains the nucleic acid binding domain (NAB). SARS-CoV NAB was shown to bind ssRNA and to unwind ATP-independent dsDNA revealing a nucleic acid chaperone like function [1,9].

Inhibitors of SARS-CoV-2 PLpro reported to date include compounds reacting with the active side Cys111, zinc conjugate inhibitors, thiopurine compounds, natural products and allosteric naphthalene inhibitors [9,10]. Naphthalene-based ligands form a large class of PLpro inhibitors with available structure-activity relationship (SAR) information. Among them is the well described allosteric inhibitor GRL-0617 which has demonstrated anti-viral activity in Vero-E6 cell infection models [11]. It has been shown to bind close to the catalytic site but not to occupy it [12]. The compound acriflavine was reported to be a potent nanomolar inhibitor of SARS-CoV-2 PLpro in enzymatic, cell based and *in-vivo* studies [13]. Using NMR and crystal structure of PLpro it was shown that two proflavine molecules occupy the substrate binding pocket. Natural products were also described as CoV PLpro inhibitors, including tanshinones, geranylated flavonoids, as well as polyphenols [1,14]. However, recent publications have arisen doubts on the mechanism of action of many of the proposed inhibitors. Ma and Wang revealed limitations of reported inhibitors by profiling them through a pipeline of enzymatic, binding and cellular activity assays, invalidating the tanshinone-family, YM155, SJB2-043, 6-thioguanine, and 6-mercaptopurine [15]. The overall homology between SARS-CoV-2 PLpro and human cysteine proteases is very low (<28% sequence identity). However, inhibitors of PLpro’s capability to cleave ubiquitin and ISG15 may impact the closest human homologues UCH-L1, USP14 and USP7 (HAUSP) [16].

## Results

### Assay development-Stability of enzymatic activity of SARS-CoV-2 PLpro

Being a multi domain protein SARS-CoV-2 nsp3 comprises multiple functions. Here we use an extended construct that comprises the PLpro and the NAB domains (named PLpro-NAB), putatively giving additional interaction points with PLpro substrate and inhibitors.

First, we evaluated two reported SARS-CoV-2 PLpro substrates: the biologically relevant substrate ISG15-AMC and as a control the Z-LRGG-AMC peptide, frequently published in PLpro biochemical screens [17]. The obtained Km values align with reported values (Fig.1) [17]. We observed no difference in Km for ISG15-AMC between the commonly used catalytic domain construct (PLpro) and the generated PLpro-NAB (S1. Fig.).

**Figure 1.**
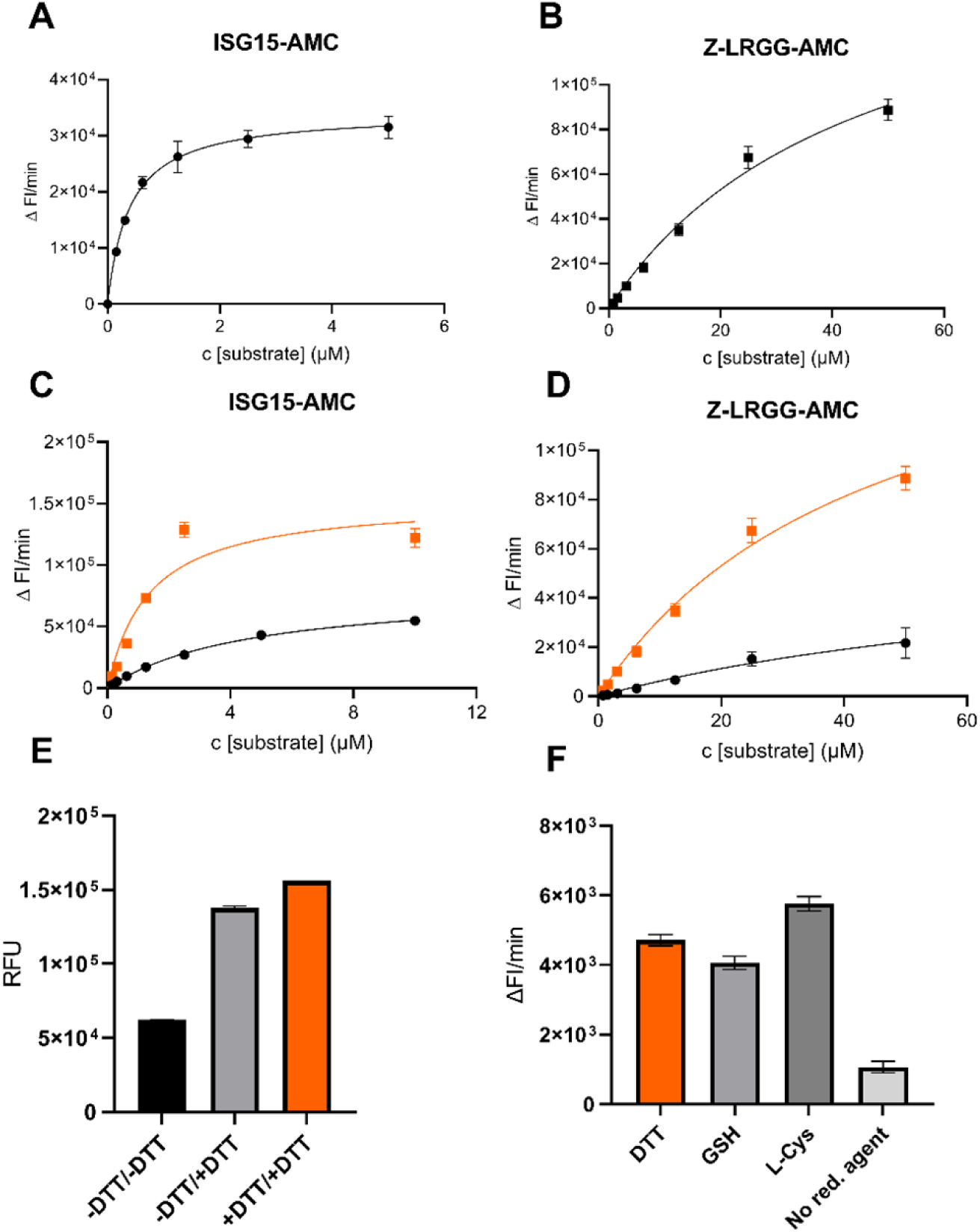
Key kinetic parameters of SARS-CoV-2 PLpro-NAB using different substrates. A: 1 nM PLpro-NAB incubated with ISG15-AMC, B: 100 nM PLpro-NAB incubated with Z-LRGG-AMC. ISG15-AMC: Vmax = 34186 /min, Km = 0.39 µM, Z-LRGG-AMC: Vmax = 173041 /min, Km = 45.2 µM. (N= 3, error bars +/-SD), PLpro activity after 30 min incubation using ISG15-AMC (C), and Z-LRGG-AMC (D) in presence and absence of DTT. **Black** line: no DTT added; **Orange** line: 1 mM DTT in assay buffer, E: Rescue effect of DTT at step of substrate addition, F: Reaction velocity different reducing agents at 1 mM in buffer (N= 3, error bars +/-SD)

To ensure protein activity over long incubation times, we evaluated enzyme activity after 30 min incubation (before addition of substrate). Increase of Km from 0.39 to 5 µM for ISG15-AMC was observed. In the presence of 1 mM DTT substrate turnover was stabilised: a five-fold smaller Km and two-fold increase of Vmax using ISG15-AMC was observed, compared to DTT-free conditions (Fig.1-C). Similar effects were seen using the Z-LRGG-AMC substrate (Fig.1-D). In addition, the loss of activity was rescued by addition of 1 mM DTT at the step of substrate addition (Fig.1-E). Different reducing agents at 1mM showed comparable results (Fig.1-F).

### Identification of new inhibitors using drug repurposing screening taking into account PLpro redox-sensitivity

Three repurposing libraries: Fraunhofer Repurposing collection (FhG), EU-Openscreen bioactives collection (EOS) and Dompe “Safe In Man” collection with 8936 compounds in total were screened against SARS-CoV-2 PLpro-NAB [18]. PR-619 and GRL-0617 were incorporated as reported positive controls for PLpro inhibition [12, 19, 20]. All screened plates passed the quality control for HTS with a Z’ > 0.5 (Mean 0.73) (S2. Fig.). Fifty-four (54) compounds, with >50 % inhibition, were selected for hit-confirmation; 50 out of 54 hits were confirmed. False positive hits, causing quenching of the AMC-signal, were removed from analysis. As DTT not only stabilizes the protein but also alters the apparent potency of reactive compounds, we evaluated inhibitor sensitivity towards reducing conditions. Hits were retested in presence of 1 mM L-cysteine, which has a lower reducing potential. Only six compounds retained their inhibition activity: CPI-169, Semapimod, SRT 1720 Sennoside A, Purpurogallin, DOM_SIM710 (3’,4’,5’,5,6,7-hexahydroxyflavone), and the positive control PR-619 (Table 1, S3. Table). Whereby SRT 1720, present two times in the test set, was not confirmed, thus discarded from further analysis. In addition, we confirmed the reference inhibitor GRL-0617 to be active in the L-cysteine buffer.

**Table 1.**
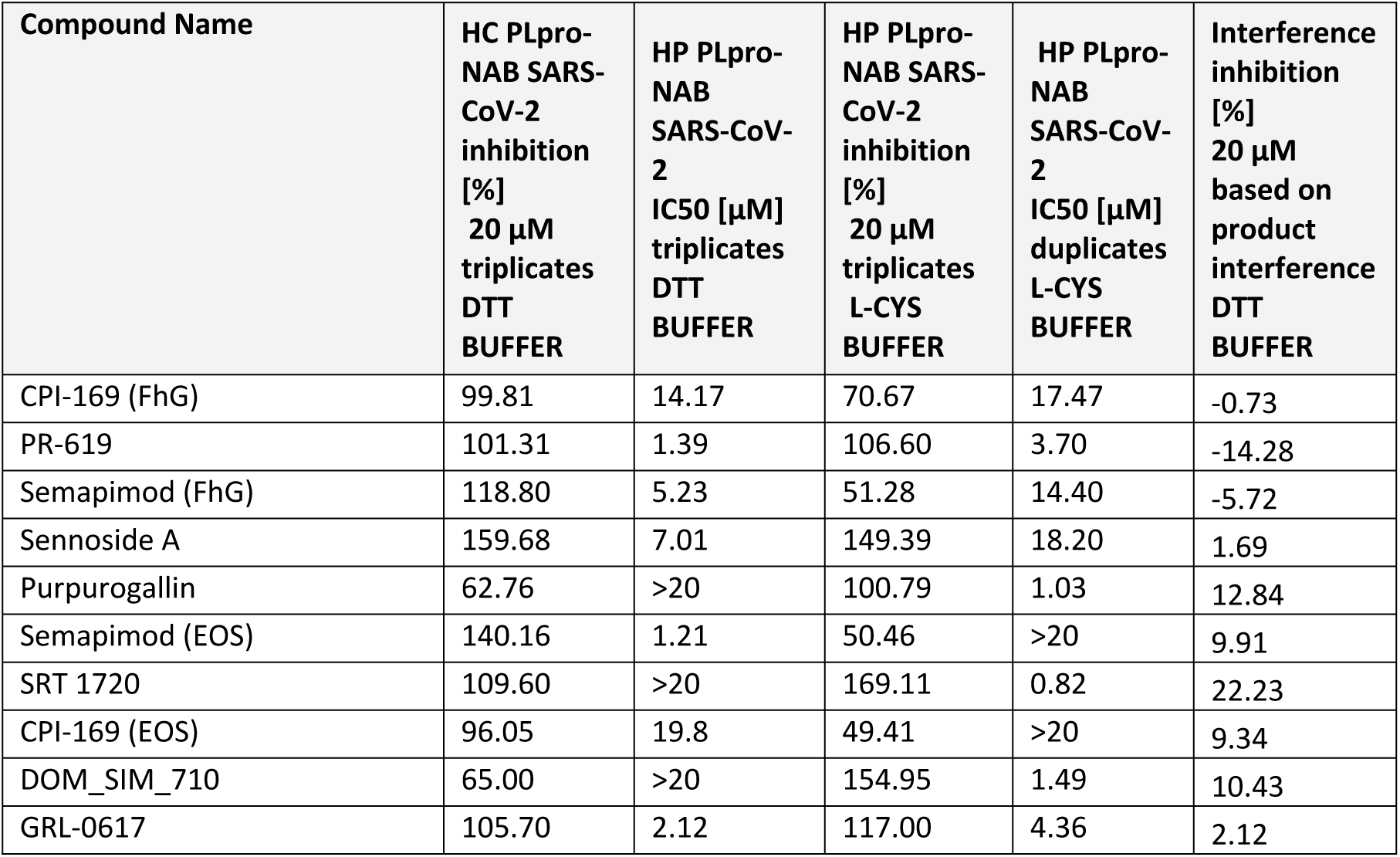
Hit compounds retaining activity in both reducing buffers with >50% inhibition. Inhibition values are normalized to activity of 20 µM PR619 (positive control) set to 100% inhibition and DMSO set to 0 % inhibition. HC-Hit Confirmation; HP-Hit Profiling.

Next, hits were analysed in dose response against SARS-CoV-2 PLpro-NAB and PLpro (Fig. 2). Walrycin B was included as the most potent hit from the quinone-like compound series, confirming its loss of activity in absence of DTT. PR-619, CPI-169, Semapimod and GRL-0617 did not show changes in potencies in different buffers. In contrast Sennoside A lost inhibitory capacity in buffer with less reducing potential (L-Cys), Purpurogallin and DOM_SIM_710 showed increased inhibitory capacity in L-cysteine buffer. In addition, we observed DOM_SIM_710 and Purpurogallin to be more potent against PLpro compared to PLpro-NAB, whereas Semapimod (FhG) was only active against PLpro-NAB, although this result was not confirmed using Semapimod from the EOS collection (data not shown).

**Figure 2.**
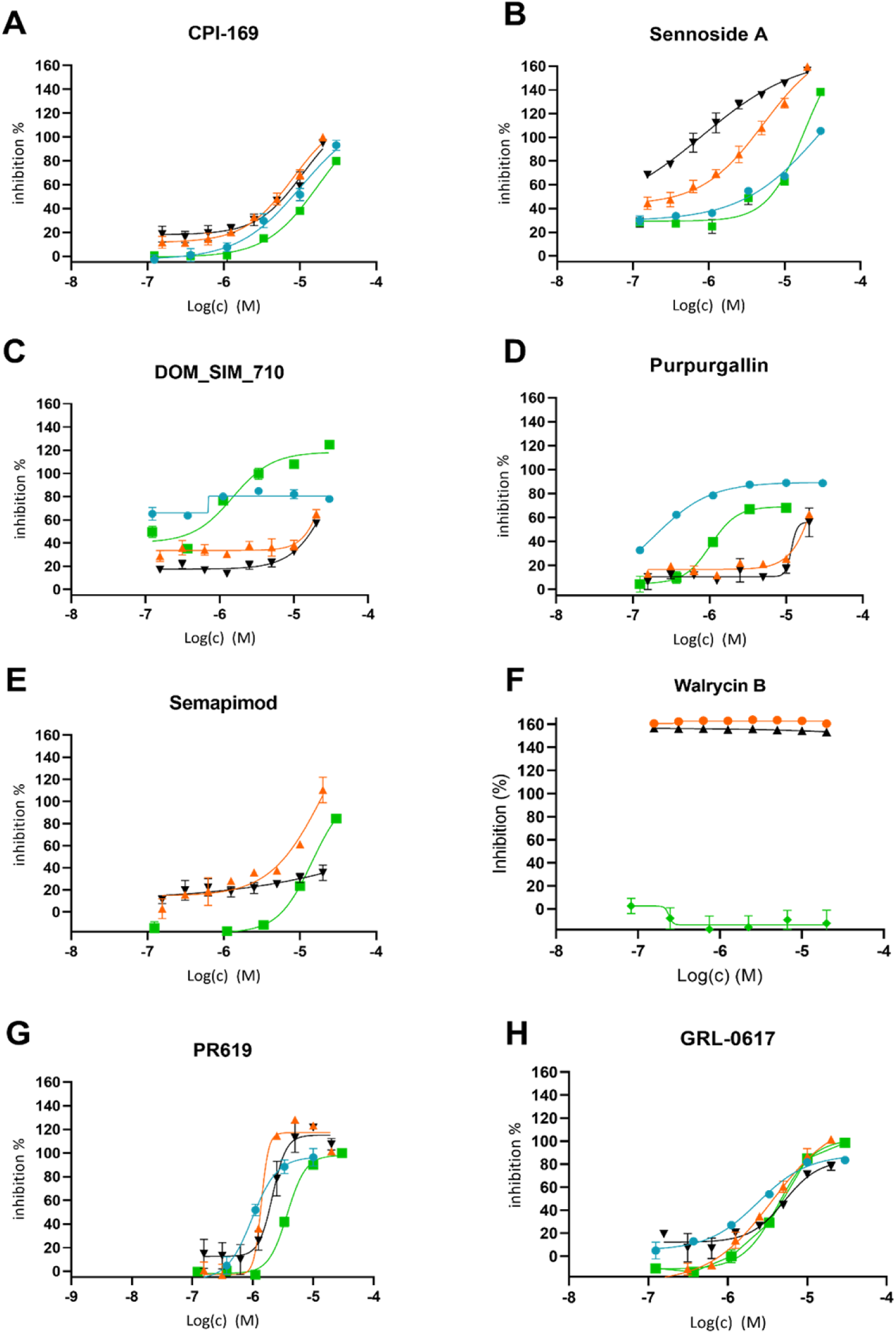
Dose response of PR-619, GRL-0617 and compounds that retained the inhibition activity against SARS-CoV-2 PLpro-NAB and PLpro under both DTT and L-Cysteine buffers. Green: PLpro-NAB/ISG15 in L-Cys Buffer; Orange: PLpro-NAB/ISG15 in DTT Buffer; Blue: PLpro/ISG15 in L-Cys Buffer Black: PLpro/ISG15 in DTT Buffer. (N= 3, error bars +/-SD). Chemical structures are shown in S5.Fig.

### Broad-spectrum activity and selectivity of identified inhibitors

PR-619, CPI-169, Semapimod, Sennoside A, Purpurogallin, DOM_SIM710, as well as GRL-0617 and Walrycin B were tested against PLpro of SARS-CoV and Mpro of SARS-CoV-2 to analyse their specificity or for broad-spectrum activity. For all tested compounds little difference was observed between PLpro of SARS-CoV-2 and SARS-CoV. Pan-inhibition of PR-619, Sennoside A and Walrycin B was observed against PLpro and Mpro SARS-CoV-2, whereby the inhibition of Mpro caused by Walrycin B was as well DTT dependent (S4. Fig). CPI-169, Semapimod, Purpurogallin, DOM-SIM_710 and GRL-0617-showed preferred inhibition of PLpro over Mpro (Fig. 3). With regards to the closest homologue proteins of SARS-CoV-2 PLpro in humans, we selected the deubiquitinase USP14 and USP7, and Cathepsin-L as representative of the cysteine protease family. All compounds, but CPI-169 and GRL-0617 showed inhibition of at least one of the human targets, with being more active compared to cathepsin-L (Fig. 3). With Walrycin B and Semapimod we observed pan-activity.

**Figure 3.**
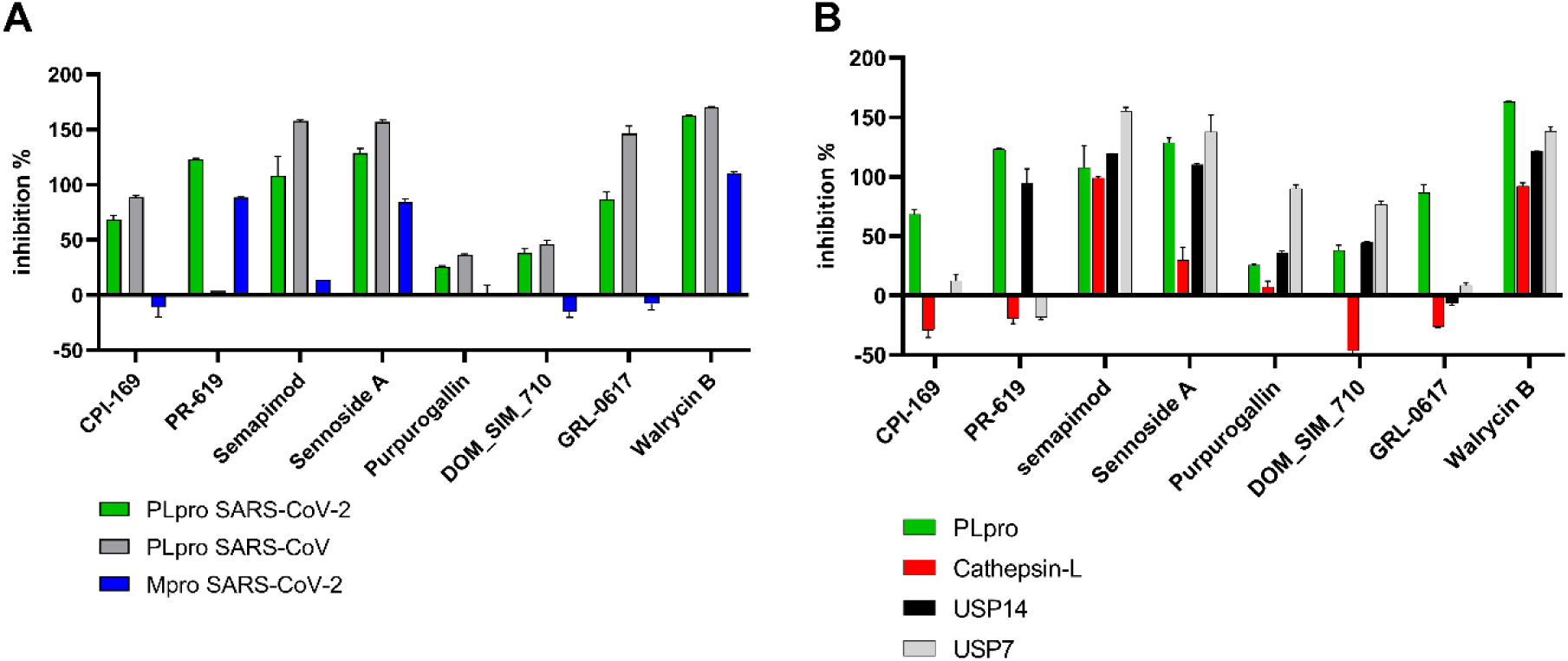
Screening for broad spectrum activity (A) and selectivity (B) of confirmed hits at 10 µM. 80% inhibition of SARS-PLpro by PR-619 at 20 µM (data not shown) (N= 3, error bars +/-SD)

### Confirmation of CPI-169 binding to PLpro

Hit profiling and selectivity tests confirmed CPI-169 as a promising inhibitor of PLpro. To verify compound binding we performed thermal-shift assays (TSA) using the PLpro-NAB construct (Fig. 4). GRL-0617 was included as control for reported binding to PLpro, and Walrycin B to evaluate the mode of inhibition for PLpro. While GRL-0617 and CPI-619 showed a concentration-dependent TSA stabilization of the PLpro-NAB, Walrycin B exhibited protein destabilization effect in presence of 1 mM DTT (Fig. 4). No influence of 1 mM L-cysteine in buffer was observed for GRL-0617 and CPI-169, while Walrycin B lost its effect on PLpro-NAB. The PLpro-NAB C111S mutant was used to verify putative activity of inhibitors towards the active site cysteine. Incubation of PLpro-NAB or PLpro-NAB C111S mutant with GRL-0617, as well as with CPI-169, resulted in increasing the Tm. The stability of the PLpro-NAB C111S was not changed in the presence of Walrycin B. Comparable results were obtained using the PLpro and PLpro C111S constructs (data not shown).

**Figure 4.**
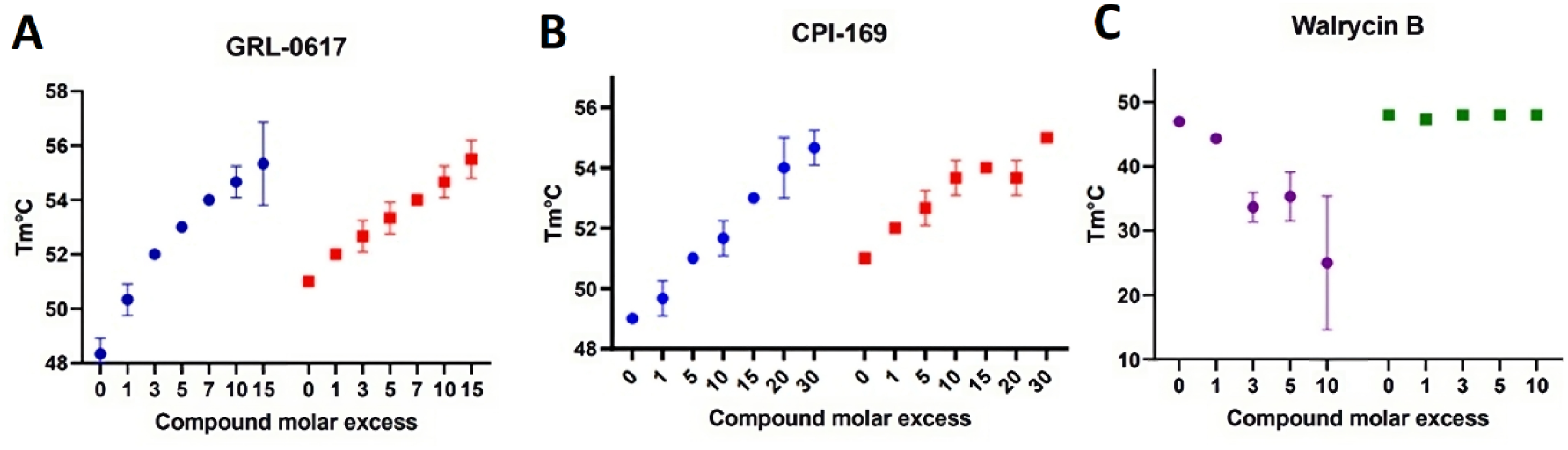
Melting temperature curve of PLpro-NAB (blue) and PLpro-NAB C111S mutant (red) measured in presence of GRL-0617 (A), CPI-169 (B) Melting temperature curve of PLpro-NAB in presence of Walrycin B and 1 mM DTT (purple) or 1 mM L-Cys (green) (C) (N= 3, error bars +/-SD)

In order to characterize modifications occurring at the catalytic cysteine due to presence of compounds, mass mapping was performed. PLpro-NAB was incubated with Walrycin B in the presence and absence of DTT. PD119507, which was previously reported as a redox-active compound generating free radicals was used as control (Table 2) [21]. In presence of DTT and the compounds, the catalytic cysteine was highly oxidized, to di-oxidized and trioxidated forms. These effects are not observed when the protein is not exposed to either compounds, or when it is treated with both compounds in the absence of a reducing agent.

**Table 2.** Percentage of modification of the catalytic cysteine for Walrycin B and PD119507 in the presence and in the absence of DTT using PLpro-NAB.

|  | Walrycin B |  | PD119507 |  |
| --- | --- | --- | --- | --- |
| DTT 1 mM | + | - | + | - |
| No-modified | 19% | 90% | 12% | 97% |
| Dioxidation | 67% | 10% | 59% | 3% |
| Trioxidation | 13% | 0% | 30% | 0% |
| Total modified | 81% | 10% | 88% | 3% |

### Docking of CPI-169 in GRL-0617 binging pocket

Due to similarities in biochemical and biophysical characteristics between CPI-169 and GRL-0617 we hypothesized a comparable binding mode. We conducted docking experiments using PLpro bound GRL-0617 as reference structure (PDB: 7JRN). We found that CPI-169 can occupy the same allosteric cavity as GRL-0617 (Fig.5). CPI-169 docking score (PLP scoring function) was in the same range of reference compound, even if it showed less potency.

**Figure 5.**
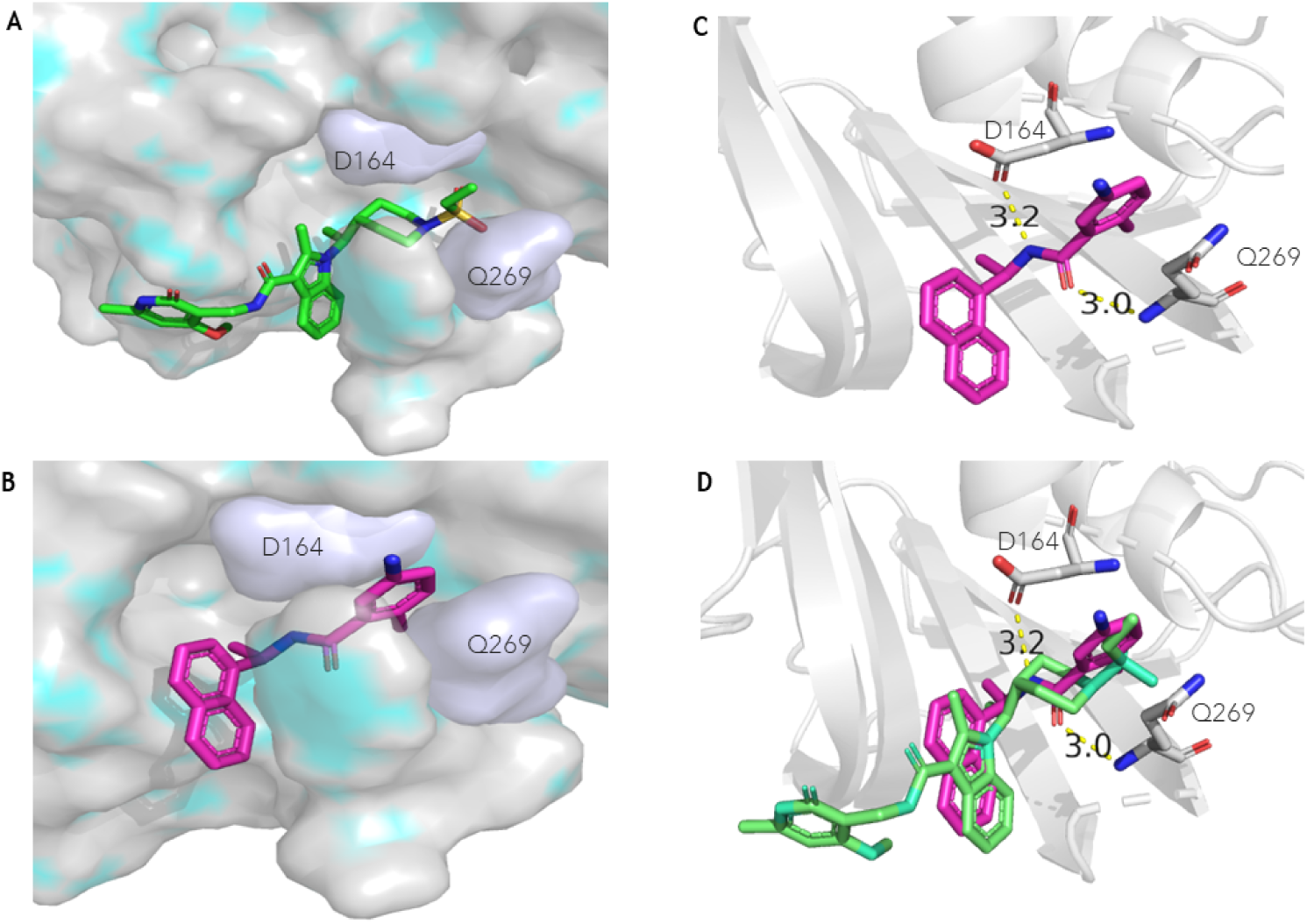
Molecular docking of CPI-169 in the allosteric pocket occupied by GRL-0617. A: Docking pose of CPI-169 in the binding pocket of PLpro. B: GRL-0617 binding pose from crystal structure (PDB 7JRN). C: GRL-0617 orientation in the binding site with highlighted H-bonds to Asp164 and Gln269. D: overlap of CPI-169 and GRL-0617 docking pose.

### Confirmation of competitive binging of CPI-169 in GRL-0619 binding pocket using STD NMR

To confirm the hypothesis of CPI-169 binding in GRL-0617 site, a competitive STD-NMR assay was performed, considering the ten times higher IC_50_ of CPI-169 compared to GRL-0617 (Fig. 6). The STD amplification factors were calculated for the methyl protons exhibiting STD signals with sufficient signal-to-noise ratio at a concentration of 0.15 mM of CPI-169 used for the competitive NMR study (S6. Fig.). Notably, tr-NOESY experiment validated the docking pose, with the only difference in the methoxide substituent of the pyridine ring, more distant from the indole moiety (S7. Table). After the addition of GRL-0167 at a concentration twice that of CPI-169, most of the amplification factors of the methyl protons of CPI-169 were reduced, while the amplification factor of H37 was significantly increased (S8. Table). The results indicate that CPI-169 is distinctly removed from the GRL-0617 binding site, while the methoxide substituent of the pyridine ring, which does not overlay with GRL-0617 in the docking model, still interacts with the outer part of the pocket, supporting the model pose of CPI-169 (Fig.5-D).

**Figure 6.**
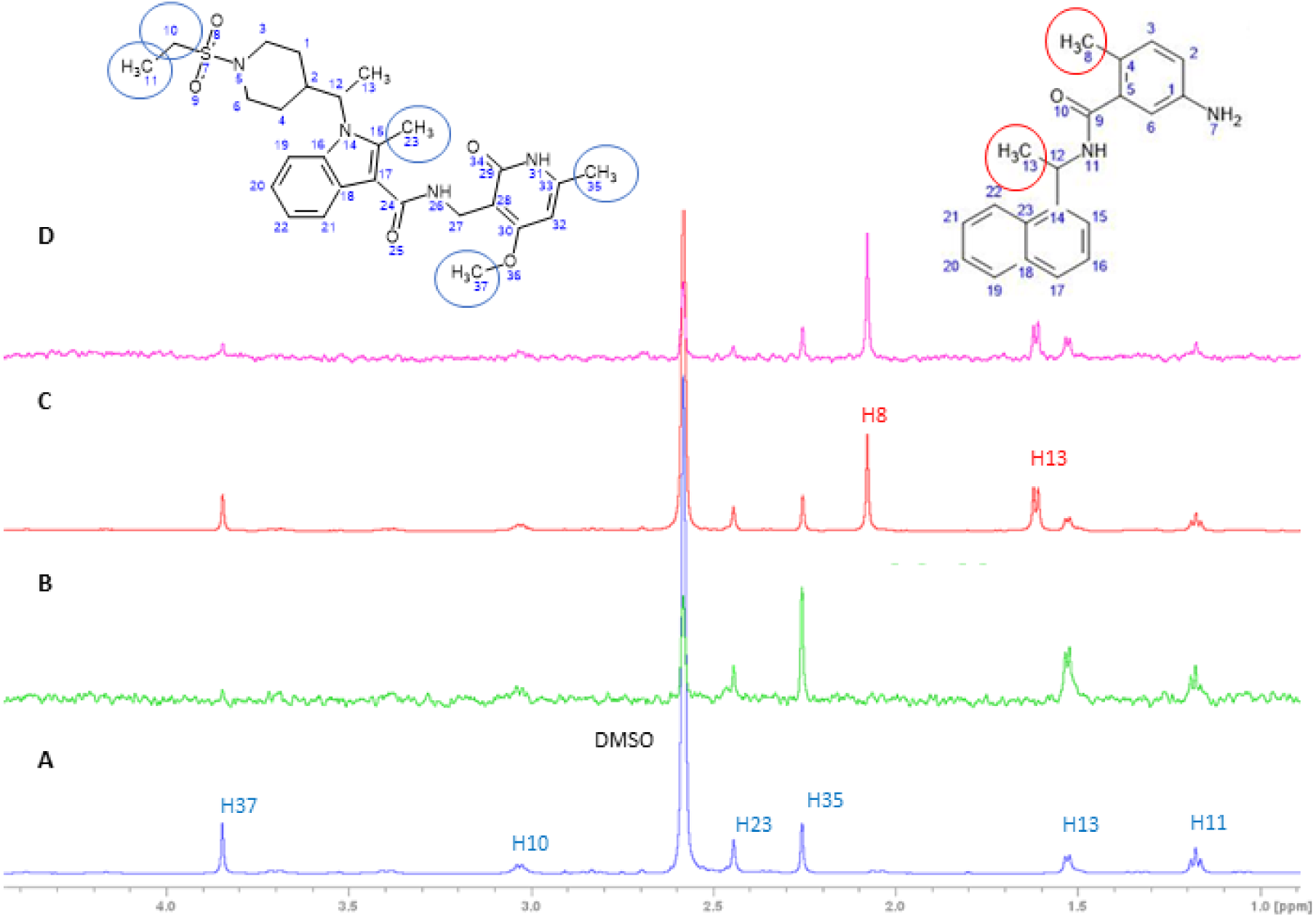
Expanded regions of 1D ^1^H STD spectra showing signals from methyl protons used to monitor competition between CPI-169 and GRL-0617. The proton signals from CPI –169 (structure on the left) and GRL-0617 (structure on the right) are labelled in blue and red, respectively. A: 1D ^1^H reference STD spectrum of 0.15 mM CPI-169 recorded at a PLpro-NAB: compound ratio of 1:100. B: 1D ^1^H STD difference spectrum of 0.15 mM CPI-169 recorded at a PLpro-NAB:CPI-169 ratio of 1:100. C: 1D ^1^H reference STD spectrum of 0.15 mM CPI-169 recorded at a PLpro-NAB: CPI-169 ratio of 1:100, in the presence of 0.3 mM GRL-0617. D: 1D ^1^H STD difference spectrum of 0.15 mM CPI-169 recorded at a PLpro-NAB:CPI-169 ratio of 1:100, in the presence of 0.3 mM GRL-0617.

## Discussion

SARS-CoV-2 PLpro is a key enzyme involved in maturation of the viral polyprotein and evasion of host immune response. Nevertheless, structural specificities, such as its active site dynamic flexibility, imply that both rational drug design and development of robust in-vitro screening assays for hit identification are challenging.

At the onset of this study, we observed high sensitivity of PLpro enzyme activity towards assay redox condition. The sensitivity of PLpro to redox agents and compounds was also reported by Arya et al., who showed that DTT (5mM) prevents aggregation of SARS-CoV-2 PLpro in biochemical assay, although with minor effect on enzymatic activity [22]. In our hands, the relatively low specific enzymatic activity of PLpro under non-reducing conditions can be reversed by the addition of DTT or L-cysteine. The most likely mechanism is the reversible oxidation of SH-groups of cysteines to sulfenic acid, especially catalytic Cys111.

To the best of our knowledge, we present here the first large-scale repurposing screen for SARS-CoV-2 PLpro using its preferred physiologically relevant substrate ISG15 in combination with the enzyme construct that includes the PLpro and the NAB domain. Primary screening in the presence of DTT gave a 0.54 % hit rate, in line with reported low hit rate of PLpro screening campaigns under similar conditions [12, 20]. However, replacing DTT with L-cysteine resulted in a loss of inhibition capacity of majority of hits. We assume that the inhibition of proteolytic activity by these compounds is due to their reaction with DTT leading to a nonspecific inhibition of PLpro. Previously, Ma et al., 2022 provided strong evidence to invalidate multiple repurposed compounds, initially reported to inhibit PLpro in biochemical assays [15]. Here, we confirm the unspecific inhibition of PLpro for many of these compounds, providing explanation related to their reactivity in strong reducing agents such as DTT. Among them, we identified a group of ortho-quinonoid and para-quinonoid derivatives, such as SF1670 and NSC663284. Previously this compound class was analysed for inhibition of CDC25, a subfamily of dual-specificity protein tyrosine phosphatases presenting a cysteine in active site. Due to activity only in presence of DTT and oxygen authors assumed a reduction to semiquinone anion radicals, producing reactive oxygen species that result in irreversible oxidation of cysteine [23]. For Walrycin B we observed similar reactivity, supporting our data in binding assays and mass mapping.

A small group of compounds: PR-619, CPI-169, Semapimod, Sennoside A, Purpurogallin, DOM_SIM710 retained their activity in both L-cysteine and DTT buffer. By analysing the specificity and broad-spectrum activity towards human target proteins USP14, USP7 and cathepsin-L, and viral SARS-CoV-2 Mpro and SARS-CoV PLpro, we revealed selective inhibition of PLpro only for CPI-169. Using TSA, and NMR we showed a similar behaviour of CPI-169 compared to GRL-0617, previously proposed by Shen et al. (preprint 2021) [24]. Both compounds bound PLpro as well as to Cys111Ser mutant of PLpro, indicating a catalytic Cysteine111 independent binding, indirectly showing possible allosteric nature of the CPI-169 inhibition. This hypothesis was proved by STD-NMR assay, through which we were able to identify the binding pocket of CPI-169 as the same as the GRL-0617 one. NMR results well match with the docking prediction and pose, highlighting weaker interactions with the PLpro with respect to GRL-0617, supporting the lower potency of this inhibitor (S6. Fig.).

## Material and Methods

### Protein expression and purification

The PLpro catalytic domain construct WT (PLpro) and the inactive mutant of the catalytic Cystein111 (PLpro C111S) cloned in *pMCSG53* vector were kindly gifted by Andrzej Joachimiak (Argonne National Laboratory, Argonne). Expression was optimized in *E.coli* BL21(DE3) by co-expressing chaperons GroEL and GroES (*pGro7* vector, TakaraBio). Cells were grown in LB medium with ampicillin (0.1 mg/mL), chloramphenicol (54 μg/mL) and L-Arabinose (0.5 mg/mL), and induced for PLpro expression with 0.5 mM IPTG and 10 μM ZnCl_2_ o/n at 20°C. PLpro WT and C111S were purified from harvested cells following the protocol reported by Osipiuk J. et al., 2021 [25]. Purified PLpro samples were stored in 20 mM Hepes, 150 mM NaCl, 1 μM ZnCl_2_, 10 mM DTT, pH 7.5, flash frozen in liquid N_2_, and preserved at –80°C.

The DNA sequence encoding for the PLpro-NAB construct (1564-2047 of nsp3), was inserted into pET24b (Novagen). The Cys111Ser mutant (PLpro^_^NAB C111S) was obtained by site-direct mutagenesis (primers 5’-GGACAACAACAGCTATCTGGCGA and 5’-GCCCATTTGATGCTGGTC). Expression was performed in BL21(DE3), grown in LB + kanamycin (50 µg/ml), and induced by 0.25 mM IPTG at 20°C for 20 hours, adding 50 μM ZnSO4. WT and C111S constructs were extracted from harvested cells by homogenization and soluble fractions loaded on a 5 ml HisTrap FF Crude column (GE Healthcare Life Sciences) equilibrated in binding buffer [20 mM Tris pH8.0, 500 mM NaCl, 10 mM Imidazole, 1 mM DTT]. PLpro fraction eluted by 0-100% gradient of elution buffer [20 mM Tris pH 8.0, 500 mM NaCl, 300 mM Imidazole, 1 mM DTT]. After negative IMAC, the TEV-cleaved proteins were purified in two steps: 1) IEX on 5 mL HiTrap Q HP (GE Healthcare Life Sciences) using as buffer A: 20 mM Bicine pH 9, 2 mM DTT, and as buffer B: buffer A + 1M NaCl; 2) SEC on Superdex 200 Hiload26/600 (GE Healthcare Life Sciences) in 20 mM Tris pH 8.0, 150 mM NaCl, 2 mM DTT. Purified fractions were concentrated to 18-25 mg/ml, aliquoted, flash frozen, and stored at –80°C till usage.

### Characterization of PLpro cysteine modifications

PLpro-NAB (7μM) was incubated with PD119507 or Walrycin B for 30 minutes at 25°C, using a molar ratio of 1:5, in the absence or presence of 1 mM DTT. The protein samples were hydrolysed with pepsin as reported in Bocedi et al., vacuum dried, and resuspended in 20 μL of HCOOH 0.2% in LC-MS Grade Water (Waters, Milford, MA) [26]. Each sample (8 μl) was analyzed by LC-MS/MS on an LTQ Orbitrap XL (Thermo Scientific, Waltham, MA) coupled to the nanoACQUITY UPLC system (Waters). Samples were concentrated onto a C18 capillary reverse-phase pre-column and fractionated onto a C18 capillary reverse-phase analytical column (250 mm, 75 μm, 1.8 μm, M-class Waters) working at a flow rate of 300 nl/min. MS/MS analyses were performed using Data-Dependent Acquisition (DDA) mode, after one full MS scan (mass range from 300 to 1800 m/z), the 5 most abundant ions were selected for the MS/MS scan events. Peptide identification analysed with Mascot software. The relative quantification of peptides containing multi-oxidised cysteine residues was carried out by employing the extracted ion current approach. The percentage of modification was calculated as the ratio of the total area of all species containing the specific modification and the total area of all the species containing the catalytic cysteine.

### Primary Assay and Screening Assay

In the PLpro-NAB primary screen, compounds, positive (20 µM PR619) and negative (100 % DMSO) controls, were transferred to 384-well assay microplates (Corning® #3820) by acoustic dispensing (Echo, Labcyte). 5 µl of SARS-CoV-2 PLpro-NAB mix were added to compound plates. Plates were sealed and incubated for 30 min at 25 °C. After addition of 5 µl ISG15-AMC (R&D Systems #UL-553) substrate, the final concentrations were: 0.15 µM substrate; 1 nM SARS-CoV-2 PLpro-NAB, 20 µM compound; and 0.2 v/v % DMSO in a total volume of 10 µL/well. The fluorescence signal was measured after 15 min of incubation with substrate (Ex/Em340/460; Envision, PerkinElmer). Assay Buffer: 50 mM Tris, 150 mM NaCl, 1 mM DTT, 0.01 % Tween20, pH 7.5. In hit confirmation and profiling, 1 mM DTT was exchanged with 1 mM L-cysteine.

Inhibition of SARS-CoV PLpro was measured using 1 nM of SARS-CoV PLpro (Biomol, #SBB-DE0024) and 0.25 µM ISG15-AMC substrate. The fluorescence signal was measured after 15 min of incubation with substrate.

### SARS-CoV-2 Mpro inhibition assay

Enzymatic activity of Mpro was measured as described previously [18]. Briefly, compounds were incubated with 60 nM Mpro for 60 min. 15 µM DABCYL-KTSAVLQ↓SGFRKM-EDANS (Bachem #4045664) substrate were added, followed by signal detection after 15 min incubation at Ex/Em= 340/460 nm using EnVision microplate reader. The assay buffer contained 20 mM Tris (pH 7.3), 100 mM NaCl and 1 mM EDTA. Zinc pyrithione at 20 µM (medchemexpress, #HY-B0572) was used as positive control, DMSO was used as solvent control.

### Cathepsin-L assay

Cathepsin-L cysteine protease activity was measured using the fluorometric cathepsin-L Inhibitor Screening Kit (BPSBioscience #79591). The assay was performed according to the manufacturer protocol, adapted to 384-well format with final volume of 10 µL. Briefly, compounds were transferred in black 384-well microplates (Corning® #3820) by acoustic dispensing (Echo, Labcyte). 5 µL of cathepsin-L are added and incubated for 30 min at 25 °C. The enzymatic reaction is initiated by addition of 5 µL substrate. Generated AFC signal is detected after 15 min at RT using Ex/Em = 360/460 nM (Envision, PerkinElmer). Final assay concentrations: cathepsin-L 0.01 ng/µL, substrate 5 µM. E-64 provided in the kit was used at 50 µM final concentration as a positive control for cathepsin-L inhibition (100 % inhibition). DMSO was used as negative control (0 % inhibition).

### USP17 and USP7 inhibition assay

5 µl/ 100 nM of USP7/USP14 (BPS bioscience, #80364) were added to assay plates containing the compounds. Plates were sealed and incubated for 30 min at 25 °, followed by addition of 5 µl/0.5 µM Ubiquitin-AMC (R&D Systems #U-550-050) substrate. The fluorescence signal was measured after 30 min of incubation (Ex/Em340/460; Envision, PerkinElmer). Inhibition by PR619 (Merck #662141)) at 200 µM was set to 100 %, DMSO to 0 % inhibition. Assay Buffer: 50 mM Tris, 150 mM NaCl, 1 mM DTT, 0.01 % Tween20, pH 7.5.

### Thermal shift assay (TSA)

TSA was performed using the PLpro-NAB (5 µM) in white 96-well plates (Bio Rad®) with a final volume of 20 µL. The assay was performed in 20 mM Tris pH 7.5, 150 mM NaCl, 1 mM DTT or L-cysteine as reducing agent. Compounds were added at increasing concentrations with final DMSO concentration of 2.5% and incubated 30 min at RT. Protein Thermal Shift Dye (Thermo Fisher Scientific®) was added at a final concentration of 0.7x from 1000x stock, emission of the dye at 560-580 nm was detected using a real-time PCR (CFX96, Bio Rad®) every 30 sec, with a temperature gradient of 2°C/min. Each analysis was executed in comparison to a negative control: buffer or compounds with dye, and a positive control: protein or protein with 2.5% of DMSO plus the fluorophore.

### NMR

All NMR experiments were recorded on a Bruker Avance Neo 600 MHz spectrometer equipped with a cryoprobe at the Slovenian NMR Centre at the National Institute of Chemistry, Ljubljana, Slovenia. Spectra were recorded at 298 K using the pulse sequences included in the Bruker TopSpin library of pulse programs. PLpro-NAB buffer was exchanged with 20 mM phosphate buffer pH 8, 50 mM NaCl with 10% deuterated water; compounds were dissolved in DMSO-d_6_. CPI-169 was tested at 0.5 mM in 6% DMSO-*d_6_*; GRL-0617 at 0.3 mM in 6% DMSO-*d_6_*. The assignment of ^1^H and ^13^C chemical shifts for CPI-169 and GRL-0617 was performed using 2D experiments, including ^1^H-^1^H TOCSY with mixing time of 0.02 s, ^1^H-^13^C HSQC, and ^1^H-^1^H tr-NOESY experiments. 1D ^1^H STD experiments [27] were performed with different concentrations while the protein:compound ratio remained 1:100.

The STD ligand epitope mapping [28] of CPI-169 was performed with 65.536 data points, 3520 scans, and a relaxation delay of 3 s using a ligand concentration of 0.5 mM. A selective on-resonance saturation of PLpro-NAB was used for 1 s at −0.772 ppm with a transmitter offset referenced to 4.699 ppm. The off-resonance irradiation was applied at 30 ppm for the reference spectrum.

The STD effect was obtained by calculating the STD amplification factors (A_STD_):

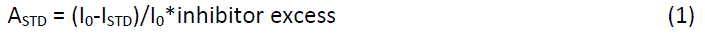

The ligand-binding epitope was represented with relative A_STD_ values normalized to the highest A_STD_ value (100%). The competitive STD experiment was performed using 0.15 mM CPI –169. Selective protein saturation was prolonged to 2 s to achieve a higher signal-to-noise ratio of STD signals at lower concentration. First, the 1D ^1^H STD spectrum was recorded at a PLpro-NAB:CPI-169 ratio of 100:1, followed by addition of GRL-0167 at GRL-0617:CPI-169 ratio of 2:1 (DMSO-d_6_ 5.5%), and the second 1D ^1^H STD spectrum was assessed, the A_STD_ of the methyl protons was calculated and compared.

The tr-NOESY spectra were acquired with a spectral width of 5882 Hz, 4096 data points in t_2_, 64 scans, 128-182 complex points in t_1_, a mixing time of 250 ms, and a relaxation delay of 1.5 s [29]. Spectra were zero-filled twice and apodized with a squared sine bell function shifted by π/2 in both dimensions.

### Docking of CPI-169

Docking of CPI-169 was performed using CCDC GOLD v2022.3.0 with PLpro bound GRL-0617 as reference structure (PDB: 7JRN). Ligand and waters were extracted and ligand pose was used for reference. Data analysis was performed among saved poses using CHEMPLP scoring function [https://doi.org/10.1006/jmbi.1996.0897].

### Data analysis

Data analysis was performed using GraphPad Prism 8 and Activitybase (IDBS). Test compound results were normalised relative to respective controls and control well outliers eliminated according to the three-sigma method. Dose response curves were fitted to 4-parameter logistic functions.

## Supporting information

**S1. Fig.**
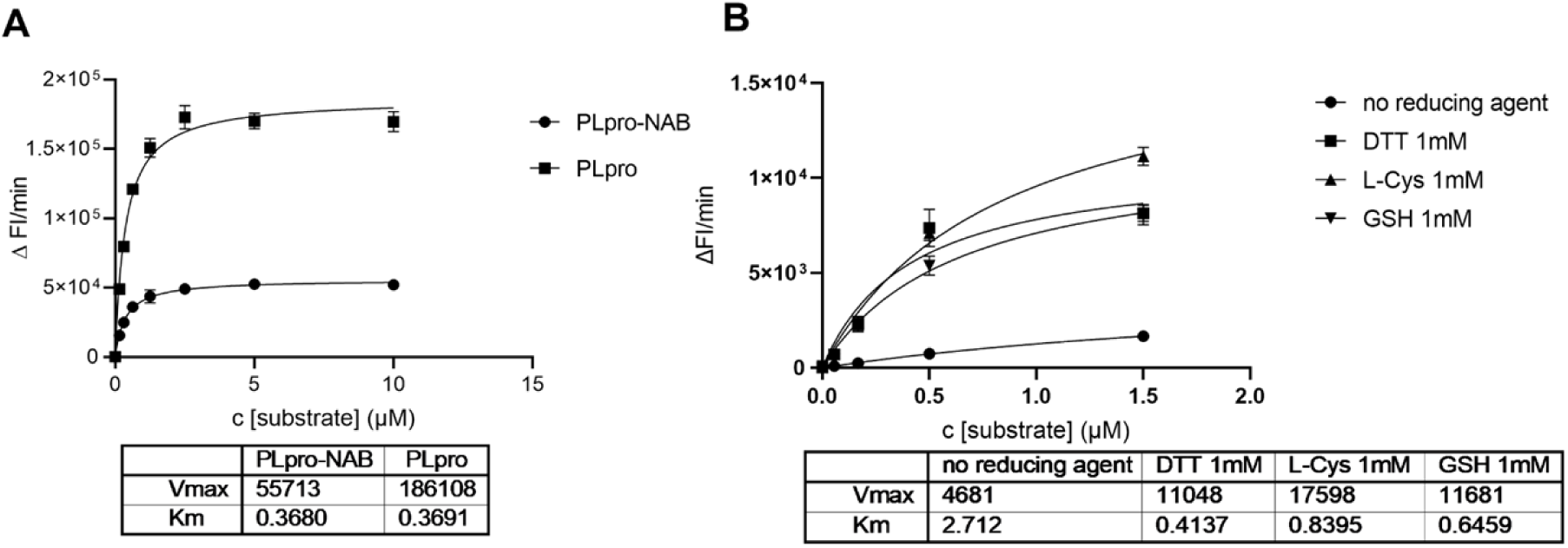
Key kinetic parameters of SARS-CoV-2 PLpro. A: Km and Vmax of PLpro containing the NAB domain (PLpro-NAB) and PLpro catalytic domain (PLpro). B: Kinetic parameters of PLpro-NAB in an assay buffer containing different reducing agents.

**S2. Fig.**
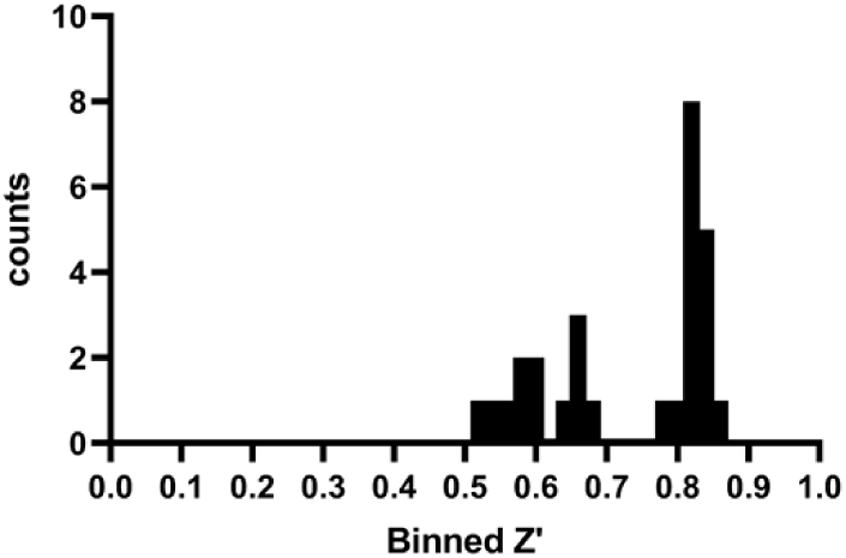
Quality control of SARS-CoV-2 PLpro-NAB primary screening. All calculated Z’ values > 0.5 showing robust assay, calculated S/B ratio value of ∼ 2.1, whereby S refers to the DMSO control and B refers to values observed using 20 µM PR-619 as inhibitor

**S3. Table.** Nominal hit compounds from PLpro-NAB primary screen with > 50 % inhibition, tested for activity under different reducing conditions. Compounds active in DTT and L-cysteine buffer are marked red, controls are marked **bold**.

| Compound Name | HC PLpro-NAB SARS-CoV-2 inhibition [%]<br>20 µM triplicates<br>DTT BUFFER | HP PLpro-NAB SARS-CoV-2 IC50 [µM]<br>triplicates<br>DTT BUFFER | HP PLpro-NAB SARS-CoV-2 inhibition [%]<br>20 µM triplicates<br>L-CYS BUFFER | HP PLpro-NAB SARS-CoV-2 IC50 [µM]<br>duplicates<br>L-CYS BUFFER | Interference inhibition [%]<br>20 µM based on product interference<br>DTT BUFFER |
| --- | --- | --- | --- | --- | --- |
| Acriflavinium Hydrochloride | 27.57 | 12.44 | -99.55 | >20 | -17.48 |
| **CPI-169 (FhG)** | 99.81 | 14.17 | 70.67 | 17.47 | -0.73 |
| Sodium-Tanshinone-ii-A-Sulfonate | 124.17 | 4.67 | -51.93 | >20 | -7.60 |
| YM-155 | 138.51 | 17.86 | -5.61 | >20 | -0.69 |
| BVT-948 | 154.88 | 0.39 | -16.22 | >20 | -137.72 |
| BYK-204165 | 137.58 | 6.17 | 7.10 | >20 | -5.54 |
| Chicago-Sky-Blue-6b | 51.02 | >20 | 3.64 | >20 | -5.59 |
| **PR-619** | **101.31** | **1.39** | **106.60** | **3.70** | **-14.28** |
| Delavirdine (mesylate) | 52.21 | >20 | -69.33 | >20 | -20.91 |
| Homidium Bromide | 149.01 | 13.69 | 18.32 | >20 | 1.40 |
| Alpha Lapachone | 146.26 | 4.43 | -6.97 | >20 | -0.29 |
| **Semapimod (FhG)** | 118.80 | 5.23 | 51.28 | 14.40 | -5.72 |
| Pyrithione Zinc | 80.42 | 20.41 | 16.01 | >20 | 3.55 |
| Propidium-Iodide | 135.55 | 0.24 | 14.90 | >20 | 0.27 |
| TAS-103 (dihydrochloride) | 154.91 | 5.94 | 17.75 | >20 | 1.87 |
| Ryuvidine | 159.90 | 0.26 | -33.69 | >20 | -2.15 |
| NSC-663284 (FhG) | 150.90 | 0.22 | 7.56 | >20 | -0.99 |
| Walrycin B (FhG) | 160.54 | 0.04 | 7.90 | >20 | 4.84 |
| Ro-08-2750 | 119.29 | 0.40 | -33.99 | >20 | 0.17 |
| Cyanocobalamin | 141.27 | 2.90 | 32.29 | >20 | 4.81 |
| **Sennoside A** | 159.68 | 7.01 | 149.39 | 18.20 | 1.69 |
| Bacitracin (Zinc) | 67.28 | 10.25 | 6.99 | >20 | -1.67 |
| 6-Hydroxy-DL-DOPA | 81.47 | >20 | 4.81 | >20 | 14.95 |
| Rose Bengal Sodium | 106.67 | >20 | -32.41 | >20 | 12.07 |
| **Purpurogallin** | 62.76 | >20 | 100.79 | 1.03 | 12.84 |
| ML113 | 74.45 | >20 | -21.50 | >20 | 10.70 |
| Juglone | 83.67 | >20 | -14.79 | >20 | 12.84 |
| PD081125 | 186.10 | 0.58 | 1.14 | >20 | 16.52 |
| PD119507 | 186.40 | 0.065 | -16.42 | >20 | 3.43 |
| ML120 | 82.78 | 3.14 | -56.07 | >20 | 3.32 |
| PT 1 | 87.07 | 9.32 | -36.09 | >20 | 9.93 |
| 9,10-Phenanthrenequinone | 180.94 | 0.504 | -40.79 | >20 | -2.46 |
| PD086277 | 180.40 | 0.807 | -30.11 | >20 | 4.30 |
| PD011561 | 49.68 | >20 | -34.83 | >20 | 13.27 |
| **Semapimod (EOS)** | 140.16 | 1.21 | 50.46 | >20 | 9.91 |
| **SRT 1720** | 109.60 | >20 | 169.11 | 0.82 | 22.23 |
| Beta-Lapachone | 175.40 | 4.71 | -37.39 | >20 | 13.40 |
| Cryptotanshinone | 40.70 | >20 | -71.36 | >20 | 3.37 |
| Dihydrotanshinone I | 80.74 | 0.92 | -176.06 | >20 | -55.01 |

| Compound Name | HC PLpro-NAB SARS-CoV-2 inhibition [%]<br>20 μM<br>triplicates<br>DTT BUFFER | HP PLpro-NAB SARS-CoV-2 IC50 [μM]<br>triplicates<br>DTT BUFFER | HP PLpro-NAB SARS-CoV-2 inhibition [%]<br>20 μM<br>triplicates<br>L-CYS<br>BUFFER | HP PLpro-NAB SARS-CoV-2 IC50 [μM]<br>duplicates<br>L-CYS<br>BUFFER | Interference inhibition [%]<br>20 μM<br>based on<br>product<br>interference<br>DTT<br>BUFFER |
| --- | --- | --- | --- | --- | --- |
| Menadione | 115.37 | 11.5 | -45.68 | >20 | -9.41 |
| Etoposide | 37.84 | >20 | -45.18 | >20 | 16.29 |
| Evans Blue | 126.18 | 8.68 | -67.82 | >20 | 6.47 |
| SF1670 | 183.62 | 0.25 | -32.73 | >20 | 9.17 |
| NSC 663284 (EOS) | 168.59 | 2.15 | -38.39 | >20 | 14.95 |
| Hexachlorophene | 109.33 | >20 | -18.33 | >20 | 15.14 |
| 3-Methyl Toxoflavin | 181.38 | 0.074 | -26.71 | >20 | 10.44 |
| Resazurin (sodium salt) | 72.88 | >20 | -36.51 | >20 | 11.91 |
| Walrycin B (EOS) | 181.14 | 0.069 | -35.90 | >20 | 8.01 |
| ML240 | 93.76 | >20 | -11.93 | >20 | 16.72 |
| CPI-169 (EOS) | 96.05 | 19.8 | 49.41 | >20 | 9.34 |
| DOM_SIM_851 | 92.36 | >20 | 24.54 | >20 | 22.72 |
| DOM_SIM_710 | 65.00 | >20 | 154.95 | 1.49 | 10.43 |
| DOM_SIM_495 | 98.55 | >20 | -39.93 | >20 | 2.55 |
| DOM_SIM_470 | 21.48 | >20 | -33.22 | >20 | 9.80 |
| **GRL-0617** | **105.70** | **2.12** | **117.00** | **4.36** | **2.12** |

**S4. Fig.**
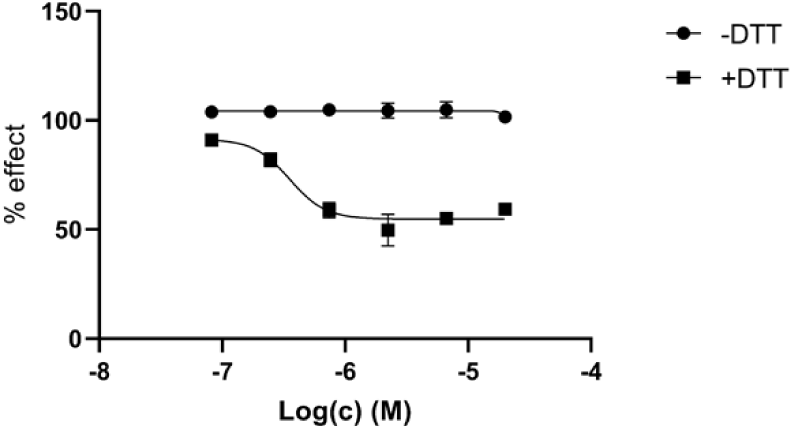
Dose dependent inhibition of SARS-CoV-2 Mpro using Walrycin B in presence and absence of DTT.

**S5. Fig.**
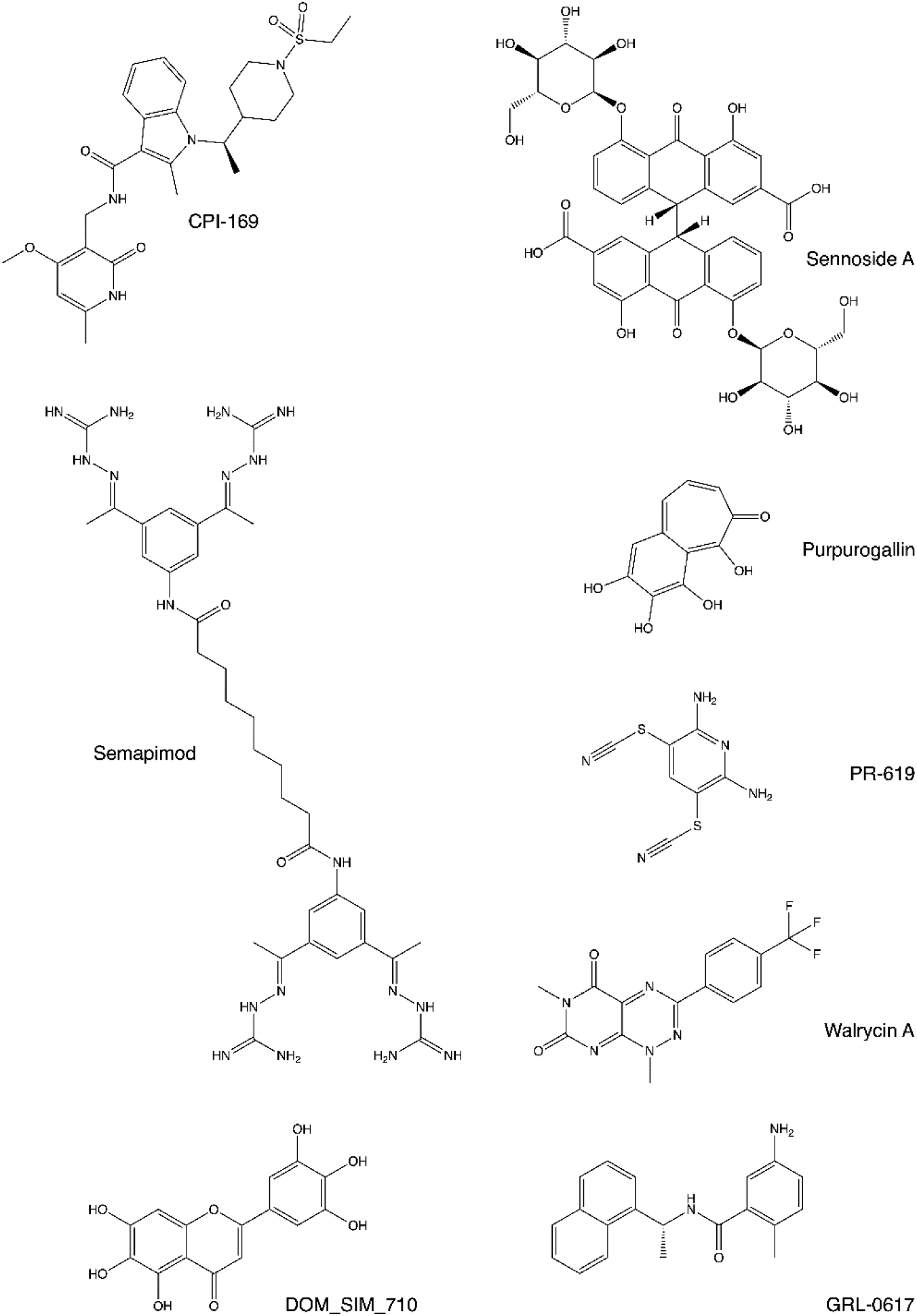
Chemical structures of PR-619, GRL-0617 and compounds that retained the inhibition activity against SARS-CoV-2 PLpro-NAB and PLpro under both DTT and L-Cysteine buffers.

**S6. Fig.**
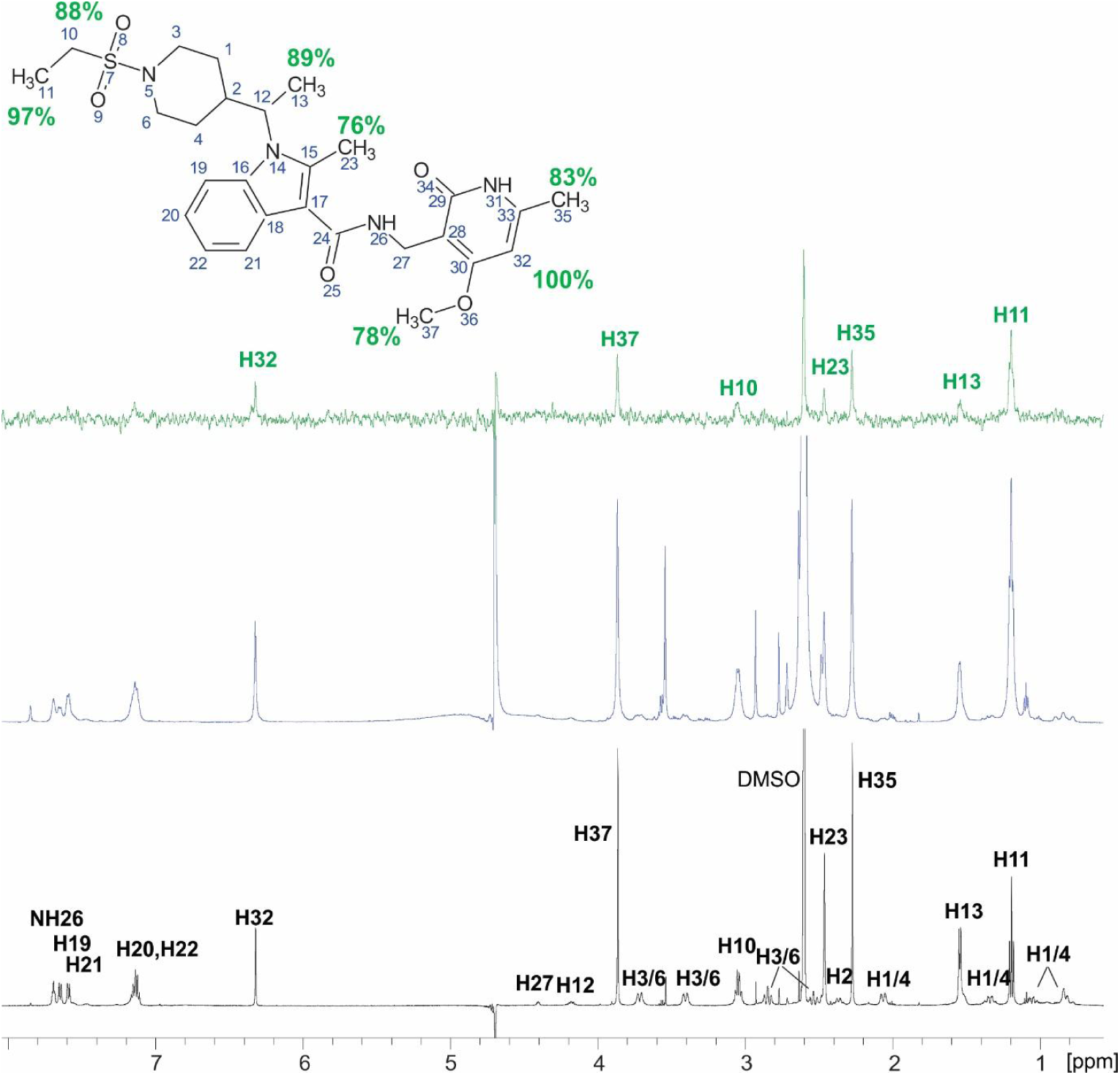
(In black) 1D ^1^H spectrum of CPI-169 showing assignment of proton chemical shifts. (In green) 1D ^1^H difference STD spectrum of CPI-169 at a PLPro-NAB:ligand ratio of 1:100. The STD amplification factors were calculated for the signals with sufficient signal-to-noise ratio marked in green and normalized to the intensity of the signal with the largest STD effect. Above, the molecular structure illustrates the proton nomenclature and the relative degrees of saturation of the individual protons. (In blue) 1D ^1^H reference (off resonance) STD spectrum.

**S7. Table.** The non-trivial NOEs of CPI-169 at a ligand-PLpro ratio of 1:100 observed in tr-NOESY spectrum and distances from the model structure of CPI-169 in complex with PLpro.

| NOE cross-peak | Intensity of NOE cross-peak | Distance from the model structure of CPI-169 in complex with PLpro (Å) |
| --- | --- | --- |
| H13-H19 | strong | 3.3 (C13-H19) |
| H13-H20,22 | weak (spin diffusion across H19) |  |
| H2-H19 | strong | 2.3 |
| H23-NH26 | weak | 4.8 (C23-NH26) |
| H37-H32 | strong | 3.0 (C37-H32) |
| H12-H19 | weak | 5.0 |
| H27-NH26 | medium | 2.6, 2.9 |
| H27-H21 | weak | 3.6, 4.3 |
|  | very weak (at decreased threshold) | 3.1 (H21-C37) |
|  | very weak (at decreased threshold) | 4.2 (H22-C37) |
| these NOEs indicate that CH<sub>3</sub>-37 has different orientation as in the model, away from H26, H21, H22 | very weak (at decreased threshold) | 2.3 (H26-C37) |
| H12-H23 | weak (spin diffusion across H3/6) |  |
| H37-H35 | weak (can be spin diffusion across H32) | 5.6 (C37-C35) |
| H3/6-H11 | weak | 4.0, 5.2 (C11-H3/6) |
| H3/6-H13 | weak (spin diffusion across H1/4) |  |
| H23-H11 | weak (spin diffusion across H12) |  |
| H23-H13 | strong | 3.9 (C23-C13) |
| H23-H1/4 | medium | 2.9, 3.7 (C23-H1/4) |

**S8. Table.** STD amplification factors of 0.15 mM CPI-619 at a PLpro/compound concentration of 1:100 (AMP) and with the addition of 0.3 mM of GRL-0617 (AMP’). AMPs of the methyl groups of CPI-169 decrease in presence of the GRL-0617, demonstrating they compete for the binding, except for the AMP of CPI-169 pyridin ring methyls, which interact with the PLpro out of the binding pocket, in line with the docking pose.

| Proton | AMP | AMP' with GRL-0167 | AMP'/AMP |
| --- | --- | --- | --- |
| H11 | 0,74 | 0.44 | 0.59 |
| H13 | 1.19 | 0.81 | 0.68 |
| H23 | 0.43 | 0.27 | 0.63 |
| H35 | 0.89 | 0.55 | 0.62 |
| H37 | 0.10 | 0.21 | 2.1 |

## Acknowledgements

The EU-OPENSCREEN bioactive compound collection was provided by the EU-OPENSCREEN ERIC (Berlin, Germany). We thank Yulia Gerhardt and Peter Maas of SPECS and Joshua Bitker for input into the selection and quality control of the Fraunhofer compound library.

## Author contributions

Maria Kuzikov: Investigation; Conceptualization; Methodology; Validation; Visualization; Formal analysis; Writing – original draft

Stefano Morasso: Resources; Investigation; Conceptualization; Methodology; Validation; Visualization; Formal analysis; Writing – original draft

Jeanette Reinshagen: Formal analysis; Data curation

Markus Wolf: Formal analysis; Data curation

Vittoria Monaco: Methodology

Flora Cozzolino: Methodology

Simona Golič Grdadolnik: Methodology

Primož Šket: Methodology

Janez Plavec: Methodology

Daniela Iaconis: Conceptualization

Vincenzo Summa: Methodology

Francesca Esposito: Methodology

Enzo Tramontano: Methodology, Conceptualization

Maria Monti: Methodology, Conceptualization

Andrea R. Beccari: Methodology, Conceptualization

Björn Windshügel: Methodology, Supervision

Philip Gribbon: Conceptualization; Funding acquisition, Supervision; Project administration

Paola Storici: Methodology, Conceptualization, Supervision

Andrea Zaliani: Conceptualization; Project administration

All authors wrote and approved the final version of the manuscript.

